# Machine learning of all *Mycobacterium tuberculosis* H37Rv RNA-seq data reveals a structured interplay between metabolism, stress response, and infection

**DOI:** 10.1101/2021.07.01.450045

**Authors:** Reo Yoo, Kevin Rychel, Saugat Poudel, Tahani Al-bulushi, Yuan Yuan, Siddharth Chauhan, Cameron Lamoureux, Bernhard O. Palsson, Anand Sastry

## Abstract

*Mycobacterium tuberculosis* is one of the most consequential human bacterial pathogens, posing a serious challenge to 21st century medicine. A key feature of its pathogenicity is its ability to adapt its transcriptional response to environmental stresses through its transcriptional regulatory network (TRN). While many studies have sought to characterize specific portions of the *M. tuberculosis* TRN, a systems level characterization and analysis of interactions among the controlling transcription factors remains to be achieved. Here, we applied an unsupervised machine learning method to modularize the *M. tuberculosis* transcriptome and describe the role of transcription factors (TFs) in the TRN. By applying Independent Component Analysis (ICA) to over 650 transcriptomic samples, we obtained 80 independently modulated gene sets known as “iModulons,” many of which correspond to known regulons. These iModulons explain 61% of the variance in the organism’s transcriptional response. We show that iModulons: 1) reveal the function of previously unknown regulons, 2) describe the transcriptional shifts that occur during environmental changes such as shifting carbon sources, oxidative stress, and virulence events, and 3) identify intrinsic clusters of transcriptional regulons that link several important metabolic systems, including lipid, cholesterol, and sulfur metabolism. This transcriptome-wide analysis of the *M. tuberculosis* TRN informs future research on effective ways to study and manipulate its transcriptional regulation, and presents a knowledge-enhanced database of all published high-quality RNA-seq data for this organism to date.

## Introduction

*Mycobacterium tuberculosis* is the leading cause of death from a single infectious agent and one of the top 10 causes of death worldwide [1]. The evolutionary success of *M. tuberculosis* is, in part, due to its adaptability to varying environments, which is largely driven by its transcriptional regulatory network (TRN) [2–4]. The TRN coordinates the expression of genes across various environmental conditions such as hypoxia, starvation, oxidative stress, and virulence. Given the global health impact of the pathogen, a deep understanding of its TRN is of fundamental importance.

One approach to TRN elucidation, successfully applied to other microorganisms, is the decomposition of a compendium of RNA-sequencing data (RNA-seq) using independent component analysis (ICA) [5–7]. ICA decomposition of transcriptomic datasets has been shown to consistently capture the underlying TRN structure in terms of sets of independently modulated genes, known as iModulons. Unlike previously defined regulons from biomolecular data describing transcription factor-DNA binding sites, iModulons are driven purely by statistical decomposition of transcriptomic data. The former is a bottom-up molecular approach focusing on individual transcriptional regulators, while the latter is a top-down global approach. ICA has already been performed on transcriptomic data compendia for *E. coli, S. aureus*, and *B. subtilis*, and has enabled a dynamic reduction in the interpretation of complex TRN responses and the discovery of new transcription factors [5–7]. While this statistical approach is useful for providing structure to the complex interactions between transcription factors (TFs), the ICA approach is heavily influenced by the volume of data available for analysis, as greater diversity of conditions in the original dataset creates a more complete iModulon structure [8].

In order to gain deeper insight into the structure and operation of *M. tuberculosis’* TRN, we performed ICA decomposition using all publicly available RNA-Seq data. We compiled 657 high quality RNA-seq expression profiles from NCBI Sequence Read Archives [9] for analysis, and extracted 80 robust iModulons. We then utilized iModulons to interpret transcriptional responses and discover molecular actors in *M. tuberculosis* transcriptional regulation by: 1) quantitatively describing the organization of the TRN and elucidating the function of new transcription factors, 2) defining transcriptional shifts that occur across changes in carbon sources, oxygen levels, and virulence states, 4) clustering various transcription factors into a core stress response stimulon. All the work described in this paper can be found at iModulonDB.org, an interactive portal for researchers to explore and download the same data used in this study

## Results

### Independent component analysis of publicly available data reveals 80 transcriptional modules for *M. tuberculosis*

In order to capture the spectrum of *M. tuberculosis’s* transcriptional response, we scraped all publicly available transcriptomic data found in NCBI’s Sequence Read Archive (SRA) and obtained 980 RNA-seq expression profiles from 53 separate studies [9] (**Figure 1a**). Each sample was processed through a standardized data processing pipeline [citation forthcoming] to assess the dataset quality and filter out poor quality profiles (See Methods, **Figure 1b**). The final dataset was composed of 657 samples, spanning various conditions that describe *M. tuberculosis*’s response to various nutrient sources, stressors, antibiotics, and virulence events. After the final dataset was obtained, a previously developed ICA algorithm was used to decompose the data into 80 robust iModulons [10] (**Figure 1b**).

**Figure 1:**
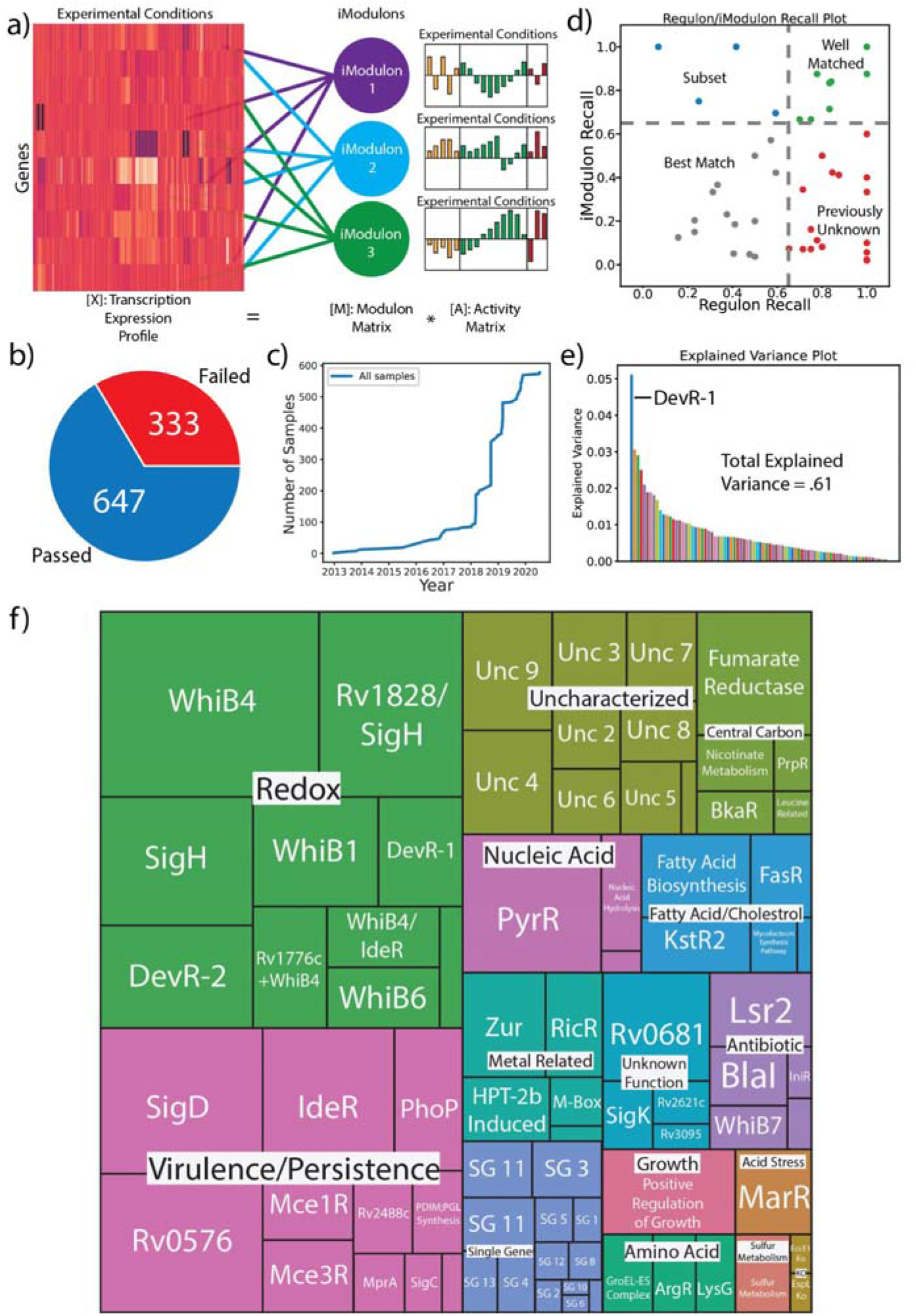
QC/QA, ICA Decomposition, and iModulon Characterization of *M. tuberculosis* RNA-seq Data from Sequence Read Archive. (a) iModulons are clusters of genes computed by decomposing RNA-Seq data into independently modulated sets (Sastry et al. 2019). (b) Percentage of samples with metadata that passed and failed the QC/QA process. The RNA-seq data and associated metadata from 980 H37Rv SRA samples were processed, and 647 samples passed all QC/QA metrics (c) A timeline of the number of high quality samples (samples that passed QC/QA) used in this study added to the Sequence Read Archive. (d) Scatter plot comparing the Regulon Recall to the iModulon Recall. iModulon Recall is defined as the number of shared genes divided by all genes in the iModulon, while Regulon Recall is defined as the number of shared genes divided by all the genes found in the regulon. iModulons in green are considered well matched, those in red contain mostly uncharacterized genes, those in blue are considered to be subsets of the regulon (i.e regulons can have multiple iModulons showing the dynamic dimensionality of the regulon), and those in grey only have a slight match. (e) Plot detailing how much explained variance is captured by each iModulon. Most iModulons capture relatively small amounts of explained variance, with the DevR-1 capturing the most variance in *M. tuberculosis*. (f) A treemap that organizes the iModulons by category. The size of each iModulon box corresponds with how many genes were found within that iModulon.

In order to provide biological interpretation of the results, iModulons were categorized by associating the set of genes in each iModulon to knowledge types, including TF binding sites, KEGG pathways, GO terms, or other associable knowledge found in the literature. An iModulon was considered associated to a particular knowledge type if there was a statistically significant (FDR < .01) overlap between the genes found in the iModulon and the knowledge type. Some iModulons were manually annotated due to shared functions of constituent genes, or presence of deleted genes (See Methods). On average, iModulons associated with transcriptional regulators could recall 66% of genes in the associated regulon, whereas literature regulons could recall 40% of the genes in the associated iModulon (**Figure 1d**)

ICA also captured the activity of each iModulon across all 657 experimental conditions, which were used to examine the response of *M. tuberculosis* to various environments. In order to minimize batch effects between the 53 studies, activity levels for each project were centered to a reference condition within the experimental subset [8]. The relative variance of iModulon activities can be mathematically related to the amount of global gene expression variation that is explained by each iModulon (**Figure 1e**). The iModulon with the highest contribution to expression variation is one of two associated with DevR, a hypoxia onset transcriptional regulator. Altogether, the 80 iModulons account for 61% of the total variance in the dataset.

After examining the mapped knowledge types and iModulon activities, each iModulon was assigned a functional category (**Figure 1f**). While most categories indicated a specific biological function, some categories indicated specific properties of the dataset. For example, the ‘Unknown Function’ category contains iModulons that have been mapped to an established TF regulon, but the function of the TF remains unclear. ‘Uncharacterized’ iModulons are those which had little overlap with known TFs or knowledge types, but still contained a significant number of genes. Finally, ‘Single Gene’ iModulons are those that track the expression of a single gene, and are treated as an artifact of the ICA decomposition [10].

We generated searchable, interactive dashboards for each iModulon and gene in our dataset, which are available at iModulonDB.org [11]. Since this genome-scale TRN covers all publicly-available high quality transcriptomic data as of August 20, 2020, other researchers are encouraged to use this site to explore the genes and regulators of interest to them.

The ICA decomposition resulted in: 1) the identification of 80 sets of independently modulated sets of genes across the entire data set (i.e., the iModulons) dramatically reducing the dimensionality of interpretation of the 3,906-gene transcriptomic response, 2) the cataloging of the activity of the iModulons under the 657 conditions, and 3) the functional annotations to the iModulons, resulting in a 61% knowledge-based description of the variation in the data set (**Figure 1e**).

### iModulons Capture the Activity of Known Transcriptional Regulators Zur and Lsr2

Two iModulons captured the action of the Zur and Lsr2 regulons, respectively. These iModulons provide a good example of how iModulons complement regulons by recapitulating expected regulator activity as well as suggesting roles under unexpected conditions. Zur is a zinc-responsive transcription factor that is activated under conditions of low zinc concentrations, such as those found in virulence conditions [12]. We found that the genes in one iModulon had significant overlap with genes in the known Zur regulon, leading us to name it the Zur iModulon (**Figure 2a**). The iModulon was found to be highly upregulated in macrophage infection conditions when compared to non-virulent controls, showing that the Zur iModulon captures the same activation conditions as the Zur TF (**Figure 2a**) [13]. Interestingly, while Zur is typically activated by zinc ions, the Zur iModulon exhibited high activities in both high and low iron concentrations. This would suggest that Zur may also be sensitive to iron ions and may play a role in the establishment of iron homeostasis together with the iron uptake regulator, IdeR. [14]

**Figure 2:**
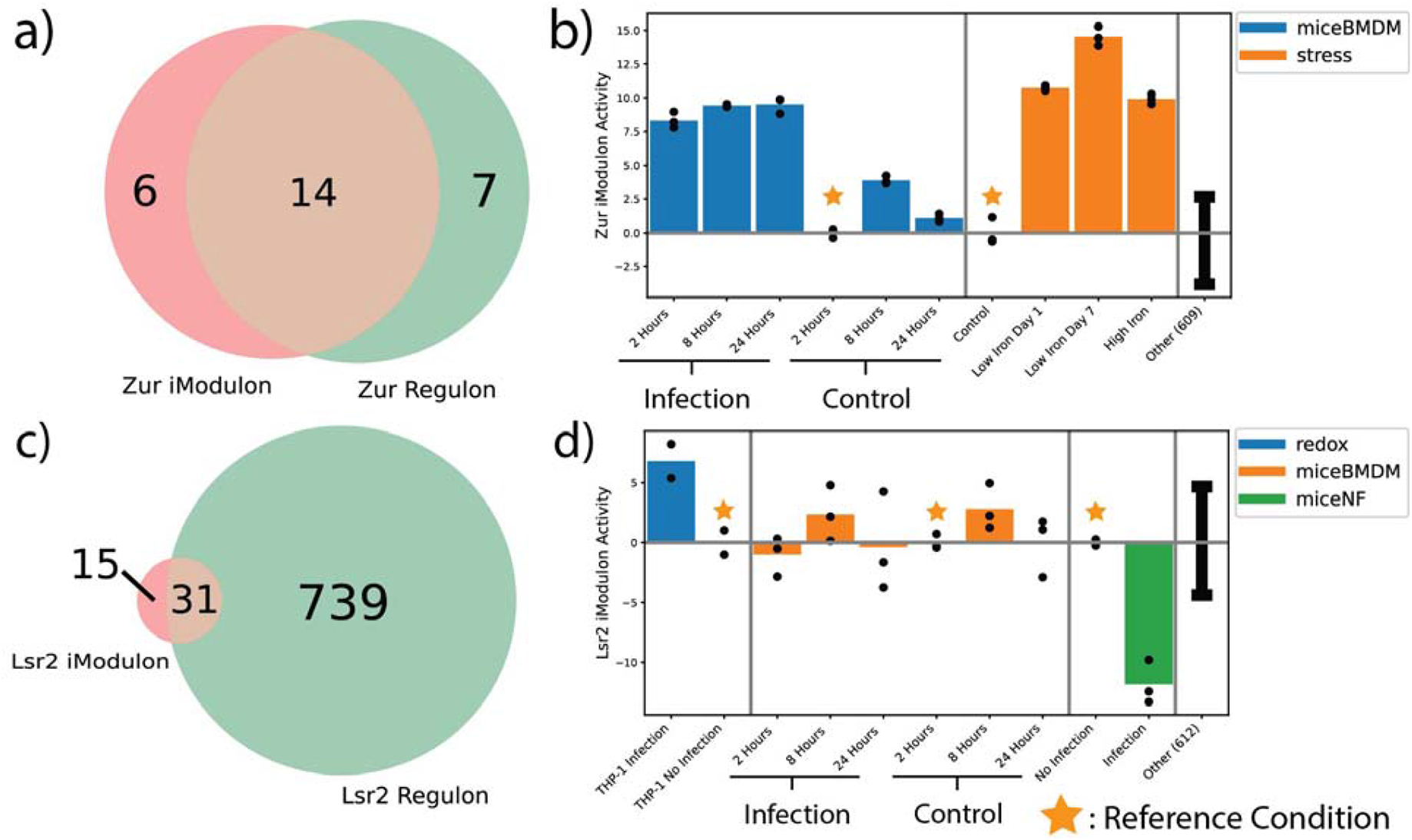
iModulons Capture Activity of Known Transcriptional Regulators Zur and Lsr2. (a) Left: Bar plot representing the activity of the Zur iModulon across virulence, high iron, and low iron conditions; Right: Venn diagram showing the genes that overlap between the established Zur regulon and the calculated iModulon (b) Left: Bar plot representing the activity of the Lsr2 iModulon across three different infection conditions (THP-1 macrophages, BMDM, and neutrophils); Right: Venn diagram showing the genes that overlap between the established Lsr2 regulon and the calculated iModulon. *For activity bar plots, error bars represent mean and standard deviation of all other samples, black dots represent the activity of each replicate for a condition, and vertical gray bars separate the samples into projects. Each project is normalized to a reference condition within that project such that the reference condition represents zero activity*.

The Lsr2 TF acts as a global transcriptional repressor that controls the expression of genes that are required for DNA organization, adaptation to oxygen levels, and chronic infections in mouse lung models [15,16]. We examined the iModulon activity under virulence virulence conditions, and found that Lsr2 may have two distinct responses depending on the cellular host (**Figure 2b**). During *in vivo* infection of murine neutrophils, the activity of the Lsr2 iModulon significantly decreased. However, during infections of mice macrophages (bone marrow derived (BMDM) or THP-1), the iModulon had significantly increased activity [13,17]. This confirmed that the iModulon activity recapitulated the prior knowledge of the TF under virulence conditions, but also suggested that Lsr2 regulation is dependent on the host cell type. We also confirmed that the iModulon was activated during hypoxia conditions, which mirrors the expected behavior of the TF [13].

### iModulons Reveal New Function of Novel Transcription Factors

Since iModulons successfully capture the structure and function of the known *M. tuberculosis* TRN, we further investigated if iModulons could discover new TF functions. Therefore, we examined the activity of the Rv0681 iModulon to determine the function of the associated TF.

Rv0681 is an uncharacterized HTH-type transcriptional regulator that has been experimentally shown to be phosphorylated by the PknH kinase [18,19]. The Rv0681 iModulon had significant overlap with the predicted Rv0681 regulon, and thus was an ideal candidate for functional discovery (**Figure 3a**) [2,4]. Upon examination, we found that a large proportion of the genes in the Rv0681 iModulon were functionally annotated to the COG classification of “lipid transport and metabolism” (**Figure 3b**). Additionally, the KstR TF, which is an important regulator for cholesterol metabolism in *M. tuberculosis*, was also found within the iModulon [20]. Given these genes, we hypothesized that Rv0681 may be involved in the regulation of lipid and cholesterol metabolism.

**Figure 3:**
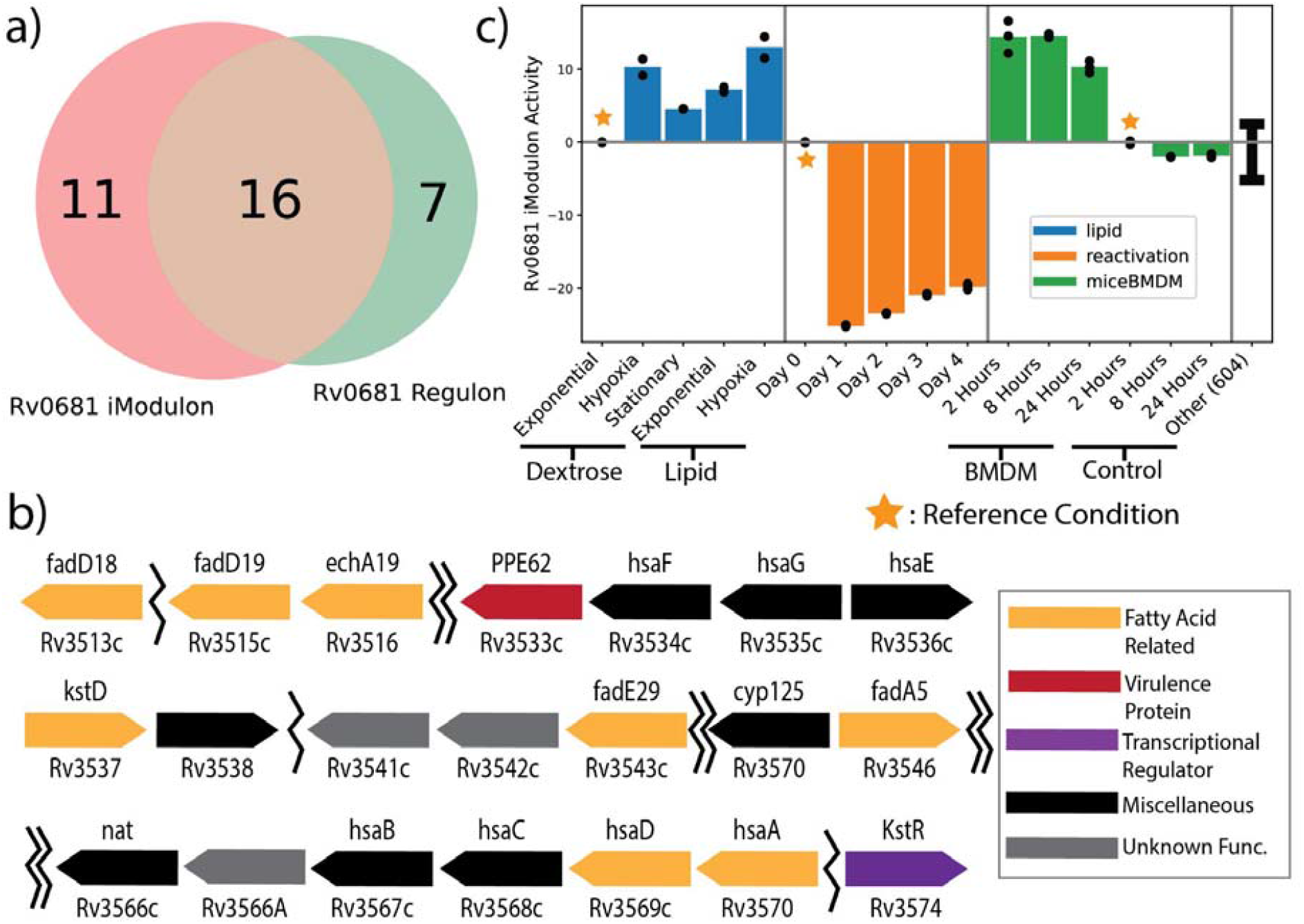
Discovery of Rv0681 as encoding a TF. (a) Venn diagram displaying the genes that overlap between the predicted Rv0681 regulon and the calculated Rv0681 iModulon. (b) Barplot displaying the activities of the Rv0618 iModulon across lipid, hypoxic reactivation, and virulence conditions. (c) A diagram that characterizes the position and function of the genes found in the Rv0681 iModulons. Many of these genes are related to fatty acids and cholesterol, including the KstR transcription factor [20]. Single jagged lines indicate a small skip between two iModulon genes (less than 10 genes), while double jagged lines indicate larger skips.

To add additional insight to the possible function of the iModulon, we examined the activity of the iModulon across three projects. In the first project, *M. tuberculosis* was grown on either dextrose or lipid-only media, during exponential phase, stationary phase, or hypoxic exposure (BioProject: PRJNA390669) [21]. We found that a switch from a dextrose media to a lipid-only media led to a significant upregulation in the Rv0681 iModulon activity, regardless of the growth phase at which the switch occurred (**Figure 3c**). This would be consistent with a function in lipid catabolism.

In a second project, *M. tuberculosis* was first induced into a persistence state via hypoxia. The bacteria was then reactivated via reaeration, and RNA-Seq was performed once a day for 4 days (BioProject: PRJNA327080) [22]. The Rv0681 iModulon had significantly decreased activity when reactivating from dormancy (**Figure 3c**), suggesting that Rv0681 is important for hypoxia and dormancy response, but is unnecessary when ample oxygen is available.

Due to the close relationship between lipids, hypoxia, and virulence, we examined a third project that tested the infection of mouse BMDM (BioProject: PRJNA478245) [3]. The iModulon was significantly upregulated during infection of the macrophage when compared to non-infection controls at all time points, confirming that the iModulon is involved with virulence as well. Altogether, we propose that Rv0681 is a transcription factor that regulates lipid metabolism (likely lipid catabolism) to promote survival in stressful conditions such as hypoxia and infection.

### Redefining the Core Lipid Response in *M. tuberculosis*

While individual iModulons can provide information about a single TF, one of their most useful functions is to simplify organism-wide transcriptional responses. Given the associated role of Rv0681 to lipid metabolism, we were interested in determining which other iModulons were activated under lipid conditions. Within the dataset, a study examined the differentially expressed genes between dextrose-and lipid-fed *M. tuberculosis* across 3 metabolic states (exponential growth, stationary phase, hypoxia) (BioProject: PRJNA390669) [21]. The study then defined a “core lipid response”, which contained genes that were found to be differentially expressed between dextrose and lipid media across all three metabolic states. This core lipid response was composed of 6 genes: Rv3161c, Rv3160c, Rv0678, Rv1217c, PPE53 and *che1* [21]. Since a core lipid response can be crucial for identifying potential targets to combat *M. tuberculosis* infections, we were interested in seeing if iModulons could be used to define a more comprehensive core lipid response utilizing the same RNA-seq data.

iModulon activities were examined between lipid and dextrose conditions, and iModulons with significant differential expression (iModulon activity change > 5 and FDR < .01) across all three metabolic states were labeled as part of the new core lipid response (**Figure 4a**). While the original study identified a core lipid response composed of only 6 genes, our analysis of the same data identified a core lipid response of four iModulons, spanning a total of 80 unique genes: Mce3R, Rv0681, Rv2488c, and Positive Regulation of Growth (PROG). The Rv0681 and Rv2488c iModulons had consistent activation across all three cell states, as Mce3R and PROG were found to have both increased and decreased activity. While the reason for this is unclear, we maintain that all four of these iModulons are important systems for *M. tuberculosis* in a lipid rich environment.

**Figure 4:**
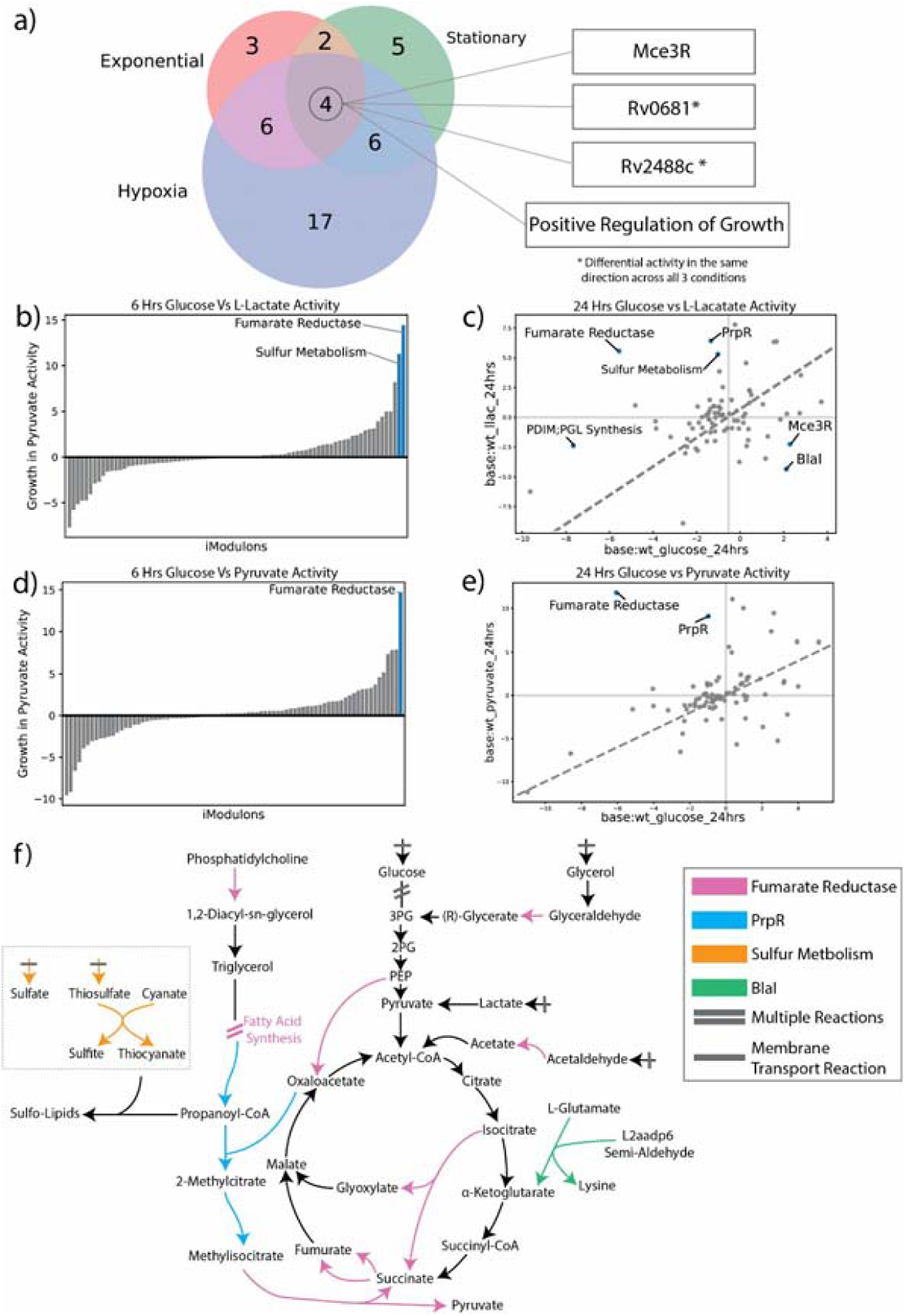
iModulons Illuminate Metabolic Shifts from Changes in Carbon Source. (a) A three-way venn displaying the differentially activated iModulons between dextrose and lipid conditions across three metabolic states (exponential, stationary, and hypoxia). The iModulons that were differentially activated across all three states represent the core lipid response. (b) A 1D DIMA plot representing the differentially activated iModulons at 6 hours between L-lactate and dextrose conditions. (c) DIMA plot representing the differentially activated iModulons at 24 hours between L-lactate and dextrose conditions. (d) A 1D DIMA plot representing the differentially activated iModulons at 6 hours between pyruvate and dextrose conditions. (e) DIMA plot representing the differentially activated iModulons at 24 hours between pyruvate and dextrose conditions. (f) A metabolic map representing the reactions controlled by differentially activated iModulons across carbon source shifts. Arrows represent reactions between metabolites, and reactions with bars represent transport from the environment. Map displays how reactions controlled by the significant iModulons are connected to one another.

Upon close examination, we found that five of the six genes previously identified as part of the core lipid response were captured by the Rv2488c iModulon, whereas *che1* was not found in any of the computed iModulons. Besides the five core lipid genes, the Rv2488c iModulon was also found to regulate the expression of various transcriptional regulators and membrane-associated proteins such as the MmpS/L efflux pump. Given the function of the clustered genes, we propose that the Rv2488c controls an essential, lipid-activated cellular defense [23]. Taken together, the results show that iModulons provide a clear definition of a core lipid response that adds to our knowledge on how *M. tuberculosis* responds to lipids.

### iModulons Elucidate Transcriptional Responses to Shifts in Carbon Sources

Given the transcriptomic response *M. tuberculosis* exhibited in a sole lipid environment, we were interested to see how the organism would respond to other carbon sources. In order to study such effects, we utilized data obtained from a study where either glucose, lactate, or pyruvate was used as a sole carbon source (BioProject: PRJNA480455) [24]. In total, the study contained six different conditions, representing the three carbon source conditions (glucose, lactate, and pyruvate) with two time points each (6 hours and 24 hours). The original study found that genes associated with the glyoxylate shunt and Krebs cycle, such as *pckA* and *icl1*, were essential and highly expressed in lactate and pyruvate conditions. To assess if iModulons could capture the upregulation of the genes highlighted in the previous findings, we created several DIMA (Differential iModulon Activity) plots to examine which iModulons were significantly changed between the glucose and the alternate carbon source (**Figure 4b-e)**. Four iModulons were of particular interest: Fumarate Reductase, Sulfur Metabolism, PrpR, and BlaI.

For cells growing on both lactate and pyruvate, the Fumarate Reductase iModulon was upregulated at all time points compared to the glucose-fed conditions. The Fumarate Reductase iModulon contains 33 genes associated with the TCA cycle and fatty acid synthesis, including *icl2, pckA*, and *fad* genes (**Figure 4b**). Many of the genes in this iModulon were also highlighted by the original study for survival in lactate and pyruvate media, which include genes that regulate the glyoxylate shunt. However, the Fumarate Reductase iModulon also captures the expression dynamics of many genes not found in the original research. These include the *fad* genes, which code for various enzymes in fatty acid synthesis, the *yrbE1* putative permeases, and the *mce1R* transcription factor, which is a vital regulator for virulence [25,26]. Many of these genes are important for maintaining lipid homeostasis, which suggests that the systems that help metabolize pyruvate and lactate are transcriptionally connected to the same systems that metabolize or synthesize lipids. Additionally, the inclusion of the virulence regulator Mce1R suggests that high levels of lactate and pyruvate may trigger the virulence response in *M. tuberculosis*.

Further evidence of a connection between alternate carbon sources, fatty acid synthesis, and virulence can be found in the significant upregulation of the Sulfur Metabolism iModulon during growth in L-lactate media. Using the iEK1008 metabolic reconstruction, we found that the Sulfur Metabolism iModulon controlled reactions for sulfate and thiosulfate import (**Figure 4f)** [27]. The presence of these sulfur-related genes in conjunction with fatty acid genes in the Fumarate Reductase iModulon strongly suggest that these pathways are involved in the synthesis of sulfolipids [27–29]. Sulfolipids are a family of sulfated acyl trehalose that aid in mycobacterium macrophage infection, and because upregulation of these pathways only occurs under lactate conditions, we propose that lactate is a cellular cue of a host cell environment.

We also found evidence of time-dependent iModulon responses during exposure to alternative carbon sources. At 24 hours, we found significant upregulation of the PrpR iModulon under both lactate and pyruvate conditions (**Figure b**,**d)**. In *M. tuberculosis*, the PrpR TF is responsible for control of the *prp* operon, which codes for several key enzymes that break down Propionyl-CoA into pyruvate and succinate, which can be used in the methylcitrate cycle to produce NADH [30]. The appearance of the PrpR iModulon at 24 hours and not at 6 hours suggests that this is a starvation response, and we hypothesize that the iModulon is activated to supplement the production of NADH and ATP from solely lactate or carbon sources. In addition, the BlaI iModulon was significantly downregulated at 24 hours, but only under lactate conditions. The BlaI TF regulates genes involved in antibiotic resistance and ATP synthesis, and the metabolic reconstruction of *M. tuberculosis* reveal that the only relevant reaction regulated by BlaI was a conversion of L-glutamate into lysine and alpha-ketoglutarate [31]. While the full function of this iModulon in response to alternative carbon sources remains unclear, its presence suggests amino acids may be involved in the synthesis of methylcitrate cycle intermediates.

### iModulon Analysis of Time-Course Data Validates Prior Models of TF Responses to Hypoxia

Hypoxia is an important signal for the TRN of *M. tuberculosis*, as the bacterium enters an altered metabolic state when exposed to reduced oxygen levels *in vitro* and *in vivo* [2,32]. While many studies have examined hypoxic response in *M. tuberculosis* by utilizing tools such as differential gene expression, here we utilized iModulons to provide additional insight on the organism’s hypoxic response.

We analyzed the important iModulons and significant activities during a hypoxia time course study in our dataset by Galagan, et al. (BioProject: PRJNA478238) [13]. During this study, the organism was exposed to changing dissolved oxygen levels, and we categorized the changes into four temporal phases: 1) Decreasing Oxygen, 2) Hypoxia Onset, 3) Stable Hypoxia, and 4) Reaeration. (**Figure 5a**). The transcriptional changes associated with hypoxia are relatively well-characterized in *M. tuberculosis*, and thus we assessed if the activities of the iModulons would recapitulate previous studies *[2]*. The study proposed a model of the *M. tuberculosis* TRN and determined that the DosR and Rv0081 TFs serve as the primary regulators for the hypoxic response, while other TFs such as Rv2034, Rv3249c, KstR, and PhoP can alter the response. In order to confirm that the iModulons recapitulate the prior model, we examined iModulons mapped to hypoxia-associated transcriptional factors and examined their activities throughout the hypoxic time course study. We found that the DevR (DosR), PhoP, KstR2, and Lsr2 iModulons had increased activity during the hypoxia time course (**Figure 5b**). The two DevR iModulons showed the highest activity during the Hypoxia Onset phase, which confirms the previous understanding that the DosR/DevR TF controls the hypoxia onset response (Figure 5a) [33].

**Figure 5:**
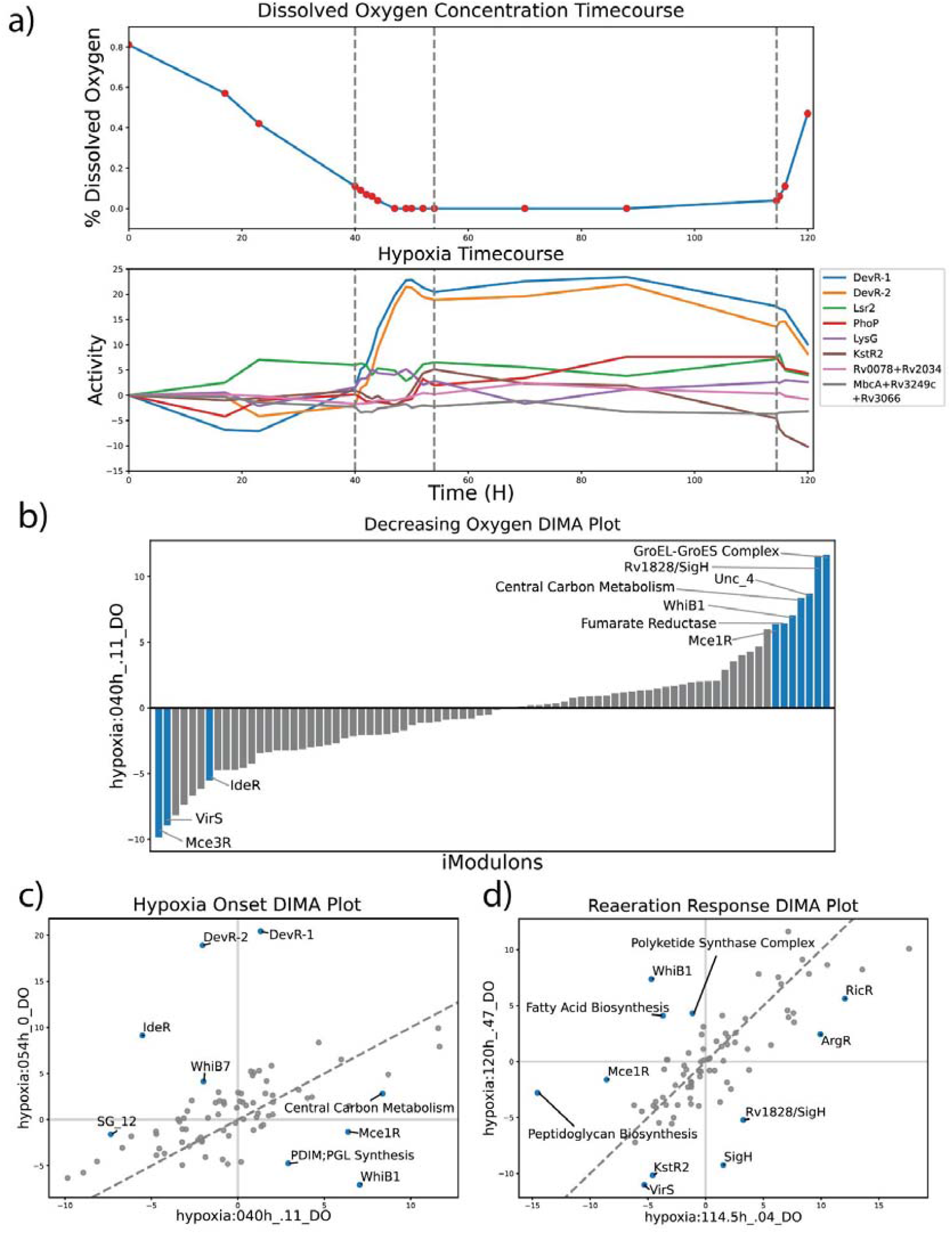
iModulons help Categorize the Phases of Hypoxia Response, including Metabolic Anticipation. (a) Time Course of *M. tuberculosis* undergoing Decreasing Oxygen, Hypoxia Onset, and Reaeration. The top plot displays the dissolved oxygen concentration in the environment, and the bottom plot displays the activities over time for iModulons controlled by TFs previously identified to be highly involved in hypoxic response. [2] The TF Rv2034 is represented by the iModulon Rv0078+Rv2034 and Rv3249c is represented by MbcA+Rv3249c+Rv3066 iModulons. (b) DIMA plots of hypoxia phases were created by comparing the iModulon activities between the first and last time point of each phase. The bar graph represents a 1D DIMA plot for the decreasing oxygen phase, since the original t=0 timepoint served as the reference condition. (c) DIMA plot for the Hypoxia Onset Phase. (d) DIMA plot for the Reaeration phase.

Additionally, the increase in activity of the Lsr2, KstR2, and PhoP iModulons also capture the known transcriptional changes associated with hypoxia, though the change in activity was relatively small compared to the DevR iModulons. Due to the lack of a KstR iModulon, KstR2 activity was examined instead as both iModulons are thought to regulate cholesterol metabolism, and thus implicate that such metabolism is important for a hypoxia response [2]. The Rv0078+Rv2034 and MbcA+Rv3249c+Rv3066 iModulons were not significantly expressed at any point in the time course.

### Different Levels of Oxygen Lead to Distinct Transcriptional States

After confirming that our iModulons are consistent with our current understanding of hypoxia, we examined the activities of the iModulons in a phase-specific manner across three of the four phases. DIMA plots were created to compare the iModulon activities from the first and last time point of each phase, and the significant iModulons were examined (**Figure 5c**,**d**). We chose not to analyze the iModulons during stable hypoxia given the limited data.

Here, we define the Decreasing Oxygen phase to represent the time when dissolved oxygen levels transition from 81% to 11%. Examination of significant iModulons during this phase reveals a three part response (**Figure 5b**). The first response is the significant increase in the production of enzymes associated with central carbon metabolism and energy production, and is captured by the Central Carbon Metabolism and Fumarate Reductase iModulons. The second response was an increased activity in cell replication systems, which was captured by the upregulation of the Rv1828/SigH, GroEL-ES complex, and WhiB1 iModulons. Rv1828/SigH contains genes that encode a wide range of proteins, including cell division proteins (SepF, FtsZ), DNA helicases (RuvA/B/C), and DNA polymerases [34]. Additionally, we found both the WhiB1 and GroEL/ES complex iModulons play a role in protein synthesis. WhiB1 also contains several genes that code for RNA polymerase, suggesting an additional role in transcription. All these iModulons have some connection with growth and replication, which suggests that cell division is an important response in *M. tuberculosis* in a decreasing oxygen environment.

The final response of the Decreasing Oxygen phase was a shift in the mammalian cell entry (Mce) proteins produced within the cell. This response is captured by increased activity in the Mce1R iModulon and a decrease in activity for the Mce3R iModulon. The Mce proteins are invasive/adhesive cell surface proteins that promote virulence in *M. tuberculosis*, and play a role in invasion of host cells [25,35,36]. Further examination of the Mce1R and Mce3R iModulons indicates that as the time course proceeds and the cell enters Hypoxia Onset and Stable Hypoxia, the activities of the two iModulons returned to their original reference state; the activity of Mce3R significantly increases while the activity of Mce1R significantly decreases. Given the close relationship between hypoxia and virulence, we theorize that Mce1 proteins help facilitate the initial stages of infections while Mce3 proteins facilitate cell entry into a dormant state.

The next phase of the hypoxia time course was the Hypoxia Onset phase, where the dissolved oxygen levels move from 11% to 0% (**Figure 5c**). Apart from the previously described activities of both DevR iModoulons, we also found that a few of the iModulons had inverted activities during Hypoxia Onset when compared to the Decreasing Oxygen phase. The Mce1R, WhiB1, and Central Carbon Metabolism iModulons showed decreased activity over the course of the Hypoxia Onset phase, indicating that high activity in these systems are either unhelpful or even detrimental to the organism during complete hypoxia. On the other hand, the IdeR iModulon moved from a decrease in activity in the prior phase to a significant increase in activity during Hypoxia Onset. Additionally, we found two new iModulons, the WhiB7 and PDIM;PGL Synthesis iModulons, with significant changes in activity during this phase. WhiB7 is a redox homeostasis transcriptional regulator that has also played a role in drug resistance [37]. The PDIM;PGL Synthesis iModulon captures genes associated with the production of phthiocerol dimycocerosate (PDIM) and phenolic glycolipids (PGL). These families of molecules have been associated with cell wall impermeability, phagocytosis, defense against nitrosative and oxidative stress and, possibly, biofilm formation [38]. The presence of both these systems during hypoxia is expected, though we did not expect PDIM;PGL Synthesis to have decreased activity during Hypoxia Onset. This would suggest that while PDIM and PGL molecules are important for oxidative stress defense, their production in a completely anaerobic environment may be detrimental to the survival of the cell.

The final phase of the hypoxia time course was the Reaeration phase (**Figure 5d**). During this phase, the cell returns to an aerobic environment as dissolved oxygen levels increase from 0% to 47%, and we found significant changes in several new iModulons. Most interesting among these are the Peptidoglycan Biosynthesis and Polyketide Synthase Complex. In *M. tuberculosis*, both polyketides and peptidoglycans are cell membrane bound molecules that play a role in cell virulence and persistence. Peptidoglycans are involved in cell growth and host response manipulation, while polyketides are essential in the formation of biofilms and are likely to improve persistence [39,40]. The increased activation of these iModulons under Reaeration suggests that *M. tuberculosis* attempts to defend itself from a possible host response during this phase. We also found that the Fatty Acid Biosynthesis iModulon had increased activity while KstR2 had decreased activity. Thus, we can conclude that under reaeration conditions, *M. tuberculosis* moves from the consumption of lipids and cholesterol to production. Taken together, the hypoxia time course and iModulons allow us to describe the complex transcriptional response that *M. tuberculosis* undergoes throughout large shifts in oxygen concentration.

### *M. tuberculosis* has Host Cell-Specific Transcriptional Responses

Due to the broad pathological impact of *M. tuberculosis*, we additionally examined the transcriptional response of *M. tuberculosis* during infection of two different host cells. In one dataset, *M. tuberculosis* was grown *in vitro* during infection of mice bone marrow-derived macrophages (BMDM), and RNA-Seq was performed at 2, 8, and 24 hours after infection (BioProject: PRJNA478245) [13]. In the other dataset, *M. tuberculosis* was grown *in vivo* in mice neutrophils, and RNA-Seq was performed at a single time point after infection (BioProject: PRJNA588440) [41]. DIMA plots were created comparing each infection condition to a control at the same time point (**Figure 6a**).

**Figure 6:**
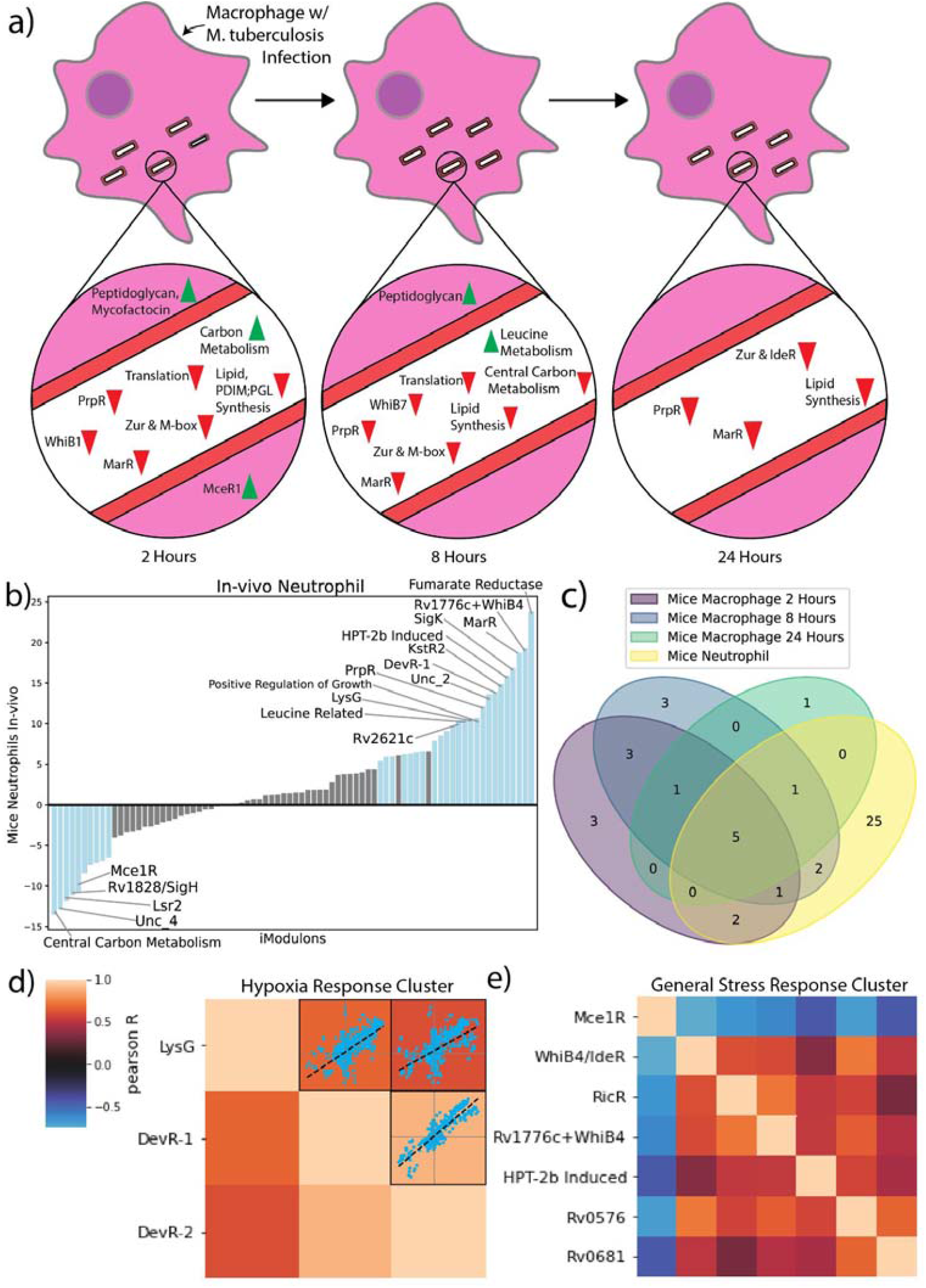
iModulon Response to Infection of Mice Macrophages and Neutrophils and Pearson R iModulon Clusters (a) A time course of the iModulon activities during infection of mice BMDM. The iModulons with differential activities at each time point are displayed as upregulated (green) or down regulated (red). Peptidoglycan, Mycofactocin, and MceR1 are displayed outside of the cell to indicate regulation of secretory pathways. **(b)** 1D DIMA plot of differential iModulons between control non-infectious condition and in-vivo infection condition. Surprisingly, the most upregulated and most downregulated iModulons both regulate different portions of central carbon metabolism, which suggests that central carbon metabolism plays a large role in infection. **(c)** A core virulence response was constructed by examining the iModulons with differential activity across all virulence conditions (3 timepoints in mice macrophage infection and 1 neutrophil condition). The core virulence response was found to consist of KstR2, MarR, PrpR, Rv0681, Uncharacterized 2, and Zur. **(d)** Hypoxia Response iModulon cluster calculated using Pearson R score and agglomerative clustering. Scatter plots that provide pairwise comparison of the activities of the iModulons across all experimental conditions is provided to indicate the relatively high correlation between these three iMoudlons. Color bar indicates pairwise Pearson R score. **(e)** General Stress Response iModulon cluster calculated from Pearson R score and agglomerative clustering.

Examination of the significant iModulons under the three time points of the mice BMDM conditions resulted in consistent patterns. Across all time points, the acid sensing MarR iModulon was found to have increased activity. MarR is a transcriptional repressor that allows *M. tuberculosis* to adapt to acidic intracellular environments [42]. In addition, we found that PrpR and lipid metabolism iModulons, along with the metal sensing Zur, M-box, and IdeR iModulons, were also consistently upregulated throughout the infection time course. All of these iModulons have been shown to play a role in either starvation or hypoxia response, indicating that residence within a macrophage requires distinct adaptations to multiple stresses [43,44].

A similar analysis of *M. tuberculosis* under *in vivo* neutrophil conditions revealed an alteredTRN response when compared to *in vitro* mice BMDM infections (**Figure 6b**). Given this difference, we investigated whether *M. tuberculosis* displayed host-cell type dependent responses. Comparison of differentially activated iModulons revealed 25 additional iModulons that were significant during infection of mice neutrophils, but not of mice BMDM. These neutrophil specific iModulons include some large regulators such as PhoP and SigK. However, we also discovered five iModulons that exhibited consistently significant activities across all experiments (KstR2, MarR, PrpR, Rv0681, Uncharacterized 2) (**Figure 6c**). All of these iModulons, with the exception of the Uncharacterized 2 iModulon, were activated in the same direction (positive activity) across the BMDM and neutrophil conditions. Overall, these results show how *M. tuberculosis* does have different transcriptional responses depending on the host cell type, but a core virulence response is required for all infection events.

### Clustering of iModulon Activities Across All Conditions Reveal Coordinated Stress Responses

By investigating the iModulons across various conditions, we noticed that certain sets of iModulons activated together. To investigate which iModulons had similar activities to one another, we clustered the iModulon activities[citation forthcoming], resulting in several clusters with biologically relevant implications. One such cluster contains the DevR-1, DevR-2, and LysG iModulons (**Figure 6d**). Given the function of DevR and the presence of the gene Rv0081 in LysG, it is clear that this cluster of iModulons captures the main hypoxic response in *M. tuberculosis* [2].

Clusters also described global responses in the *M. tuberculosis* TRN, as shown by the General Stress Response Cluster (Figure 6e). This cluster contained virulence iModulons such as Mce1R, metal related iModulons like RicR, and lipid metabolism iModulons such as Rv0681. We found that while six of the iModulons within the cluster were positively correlated with each other, Mce1R was found to be negatively correlated with the others. To help visualize which systems were controlled by this cluster, we mapped the genes within each associated iModulon to known pathways using a metabolic reconstruction [27]. The reactions encoded by the iModulons in the cluster linked cholesterol-catabolism pathways to propionyl-CoA biosynthesis. Propionyl-CoA is an important precursor to sulfolipids, and we found that the General Stress Response Cluster also controls pathways associated with sulfur import and the formation of sulfolipids. The cluster also controls the production of mce1 proteins, the type 1 NADH-dehydrogenase, and metal sensing systems. Type 1 NADH-dehydrogenase is known to produce ROS species and increase oxidative stress, while metal sensing systems such as those encoded by RicR are important for protection against oxidative stress [45,46]. Given the function of these genes, we propose that this cluster represents a general stress response in *M. tuberculosis*, most likely related to intra-host survival. Though the General Stress Response Cluster represents a commonly co-transcribed set of iModulons, each one is still independently modulated; there are instances where one part of the cluster is not needed and its iModulon’s activity diverges from the rest. This example demonstrates that iModulon clustering can create a complex, hierarchical understanding of the TRN.

## Discussion

Here, we utilized ICA to decompose 657 distinct RNA-Seq profiles of *M. tuberculosis* into 80 independently modulated sets of genes, termed iModulons. Many of these iModulons correspond to important transcription factors in the organism. Using these iModulons, we 1) revealed the function of previously unknown regulons, 2) described the transcriptional shifts that occurred during environmental changes such as carbon source shifts, oxidative stress induction, and infections, and 3) demonstrated the presence of large clusters of transcriptional regulons that link several important metabolic systems, including lipid, cholesterol, and sulfur metabolism.

In contrast to standard molecular biology approaches, which build knowledge of the transcriptome in a bottom-up fashion by observing biomolecular binding events, ICA decomposition builds understanding in a top-down fashion by extracting signals from large datasets. ICA can thus extract coherent regulation without knowledge of an associated TF, leading to the discovery of unknown regulons and regulon functions, and ultimately elucidating a more complete TRN.

We demonstrated that iModulons are effective at providing mechanistic insights into complex transcriptional changes in the TRN. Lipid metabolism, hypoxia protection, and host cell responses are all vital factors in the success of *M. tuberculosis* as a pathogen. iModulons provided a clear model of the transcriptome changes occurring under these conditions. Such mechanistic breakdown of TRN responses can also be applied to antibiotic exposure and resistance, providing further understanding on how to combat *M. tuberculosis* infections.

Additionally, the *M. tuberculosis* iModulons were clustered together based on their activities, revealing system-wide stress responses. These clusters suggest the presence of transcriptional stimulons, or clusters of genes that all respond to the same stimulus. Such stimulons, especially ones that respond to antibiotics, can also provide better understanding on ways to combat the pathogen.

All results presented in this manuscript are reproducible at [add github]. In addition, we have provided an interactive portal for any researcher to investigate iModulon structure of *M. tuberculosis*, the iModulon activities, and the original gene expression compendium at iModulonDB.com. Given the size of the dataset, this data still has potential to reveal new insights into the function of uncharacterized transcription factors and the TRN behavior of *M. tuberculosis* under different conditions. With the elucidation of the iModulon structure of *M. tuberculosis*, it is now possible to decompose other Mycobacterium strains that are used to study H37Rv, such as H37Ra and *M. Smegmatis* [cite]. Such analysis can reveal the differences in the transcriptomic organization of these related strains and provide further information of the. In summary, the ICA decomposition on mycobacterium and other bacterial transcriptomics data is still rich with new discoveries.

## Supporting information

Supplemental Table 1 - Dataset Citations

Supplemental Table 2 - TRN Citations

Supplemental Table 3 - Lipid Core

Supplemental Table 4 - Carbon Source Shift

Supplemental Table 5 - Hypoxia Timecourse

Supplemental Table 6 - Virulence

Supplemental Figure 1 - Pearson Cluster Map

Supplemental Figure 2 - Core Stress Response Metabolic Map

## Data Availability

The iModulons composition, activities, and the code used to generate figures and results are available on Github (https://github.com/Reosu/modulome_mtb). Detailed, curated dashboards for each iModulon and gene can be searched or browsed on iModulonDB.org under the “*M. tuberculosis* Modulome” dataset (https://imodulondb.org/).

## Methods

The functions used in this study and description of the methods for compiling and processing RNA-Seq data, running ICA, and computing iModulon enrichments were adapted from the Pymodulon methods paper from Sastry et al. [citation forthcoming].

### Compiling all public transcriptomics data

Using the script from Sastry et al. [citation forthcoming], (https://github.com/avsastry/modulome_workflow/tree/main/download_metadata), we found all RNA-seq data for *M. tuberculosis* on NCBI SRA as of August 20, 2020. We manually selected samples that used the strain *M. tuberculosis* H37Rv.

### Processing prokaryotic RNA-seq data

To process the complete *M. tuberculosis* RNA-seq dataset, we used Amazon Web Services (AWS) Batch to run a Nextflow pipeline.

The first step in the pipeline is to download the raw FASTQ files from NCBI using fasterq-dump (https://github.com/ncbi/sra-tools/wiki/HowTo:-fasterq-dump). Next, read trimming is performed using Trim Galore (https://www.bioinformatics.babraham.ac.uk/projects/trim_galore/) with the default options, followed by FastQC (http://www.bioinformatics.babraham.ac.uk/projects/fastqc/) on the trimmed reads. Next, reads are aligned to the genome using Bowtie [47]. The read direction is inferred using RSEQC [48] before generating read counts using featureCounts [49]. Finally, all quality control metrics are compiled using MultiQC [50] and the final expression dataset is reported in units of log-transformed Transcripts per Million (log-TPM).

### Quality Control and Data Normalization

To guarantee a high quality expression dataset for *M. tuberculosis*, data that failed any of the following four FASTQC metrics were discarded: per base sequence quality, per sequence quality scores, per base n content, and adapter content. Samples that contained under 500,000 reads mapped to coding sequences were also discarded. Hierarchical clustering was used to identify samples that did not conform to a typical expression profile..

Manual metadata curation was performed on the data that passed the first four quality control steps. Information including the strain description, base media, carbon source, treatments, and temperature were pulled from the literature. Each project was assigned a short unique name, and each condition within a project was also assigned a unique name to identify biological and technical replicates. After curation, samples were discarded if (a) metadata was not available, (b) samples did not have replicates, or (c) the Pearson R correlation between replicates was below 0.95. Finally, the log-TPM data within each project was centered to a project-specific reference condition.

### Computing the optimal number of robust Independent Components

To compute the optimal independent components, an extension of ICA was performed on the RNA-seq dataset as described in McConn et al. [10].

Briefly, the scikit-learn (v0.23.2) [51] implementation of FastICA [52] was executed 100 times with random seeds and a convergence tolerance of 10^−7^. The resulting independent components (ICs) were clustered using DBSCAN [53] to identify robust ICs, using an epsilon of 0.1 and minimum cluster seed size of 50. To account for identical with opposite signs, the following distance metric was used for computing the distance matrix:

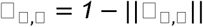

where ρ_*x,y*_ is the Pearson correlation between components *x* and *y*. The final robust ICs were defined as the centroids of the cluster.

Since the number of dimensions selected in ICA can alter the results, we applied the above procedure to the *M. tuberculosis* dataset multiple times, ranging the number of dimensions from 10 to 320 with a step size of 20. To identify the optimal dimensionality, we compared the number of ICs with single genes to the number of ICs that were correlated (Pearson R > 0.7) with the ICs in the largest dimension. We selected the number of dimensions where the number of non-single gene ICs was equal to the number of final components in that dimension.

### Compiling gene annotations

The gene annotation pipeline can be found at https://github.com/SBRG/pymodulon/blob/master/docs/tutorials/creating_the_gene_table.ipynb. Gene annotations were pulled from AL009126.3. Additionally, KEGG [54] and Cluster of Orthologous Groups (COG) information were obtained using EggNOG mapper [55]. Uniprot IDs were obtained using the Uniprot ID mapper [56], and operon information was obtained from Biocyc [57]. Gene ontology (GO) annotations were obtained from AmiGO2 [58]. The known transcriptional regulatory network was obtained primarily from the Galagan predicted binding database and MTB Network portal databases. [2,4]

### Computing iModulon enrichments

iModulon enrichments against known regulons were computed using Fisher’s Exact Test, with the false discovery rate (FDR) controlled at 10^−5^ using the Benjamini-Hochberg correction. Fisher’s Exact Test was used to identify GO and KEGG annotations as well, with an FDR < 0.01.

Additional functions for gene set enrichment analysis are located in the *enrichment* package, including a generalized gene set enrichment function and an implementation of the Bonferroni-Hochberg false discovery rate (FDR).

### Calculating Differentially Expressed iModulons Across Conditions

The difference in activity of iModulons were compared across relevant conditions and significantly changed iModulons were calculated utilizing a lognormal probability distribution. For each comparison, we computed the absolute difference in the mean iModulon activity and compared it to an iModulon’s log-normal distribution (calculated between biological replicates). P-value statistics was obtained for a given pair of conditions across all iModulons and a FDR was calculated. iModulon changes were considered significant if the difference was greater than 5 and FDR < .01.

DIMA scatter plots plot the activities of iModulons under one condition versus another, and allow for the visualization of significantly changed iModulons. 1D DIMA plots plot iModulons under one condition to a reference condition. Reference conditions have been normalized to have 0 activity across all iModulons, and thus a bar plot is used instead of a scatter plot.

### Calculating iModulon Activity Clusters

The activities of iModulons were clustered using a Seaborn clustermap. [59] Pearson R correlation was used as a distance metric, and pairwise distances for each iModulon were calculated. After creation of the clustermap, the scikit-learn agglomerative clustering function was performed on the clustermap [51].

Optimal cluster sizes were obtained by computing the varying the threshold statistic for agglomerative clustering and finding the optimal silhouette score. Once iModulons clusters were calculated, clusters that had above average Pearson R correlation between iModulons were manually inspected to determine physiological function.

### Generating iModulonDB Dashboards

iModulonDB dashboards were generated using the PyModulon package [citation forthcoming], [11]]; the pipeline for doing so can be found at https://pymodulon.readthedocs.io/en/latest/tutorials/creating_an_imodulondb_dashboard.html. Where applicable, we provide links to gene information in Mycobrowser [60].

## Acknowledgements

We thank Erol Kavvas, Nick Dillon, and Amitesh Anand for helpful discussions and biological insights. This research used resources of the National Energy Research Scientific Computing Center, a DOE Office of Science User Facility supported by the Office of Science of the U.S. Department of Energy under Contract No. DE-AC02-05CH11231.

## Notes

https://github.com/Reosu/modulome_mtb

https://imodulondb.org/index.html

### Competing Interest Statement

The authors have declared no competing interest.

