## Supplementary figures and images for "Machine learning of all *Mycobacterium tuberculosis* H37Rv RNA-seq data reveals a structured interplay between metabolism, stress response, and infection"

### Supplemental Figure 1 - Pearson Cluster Map

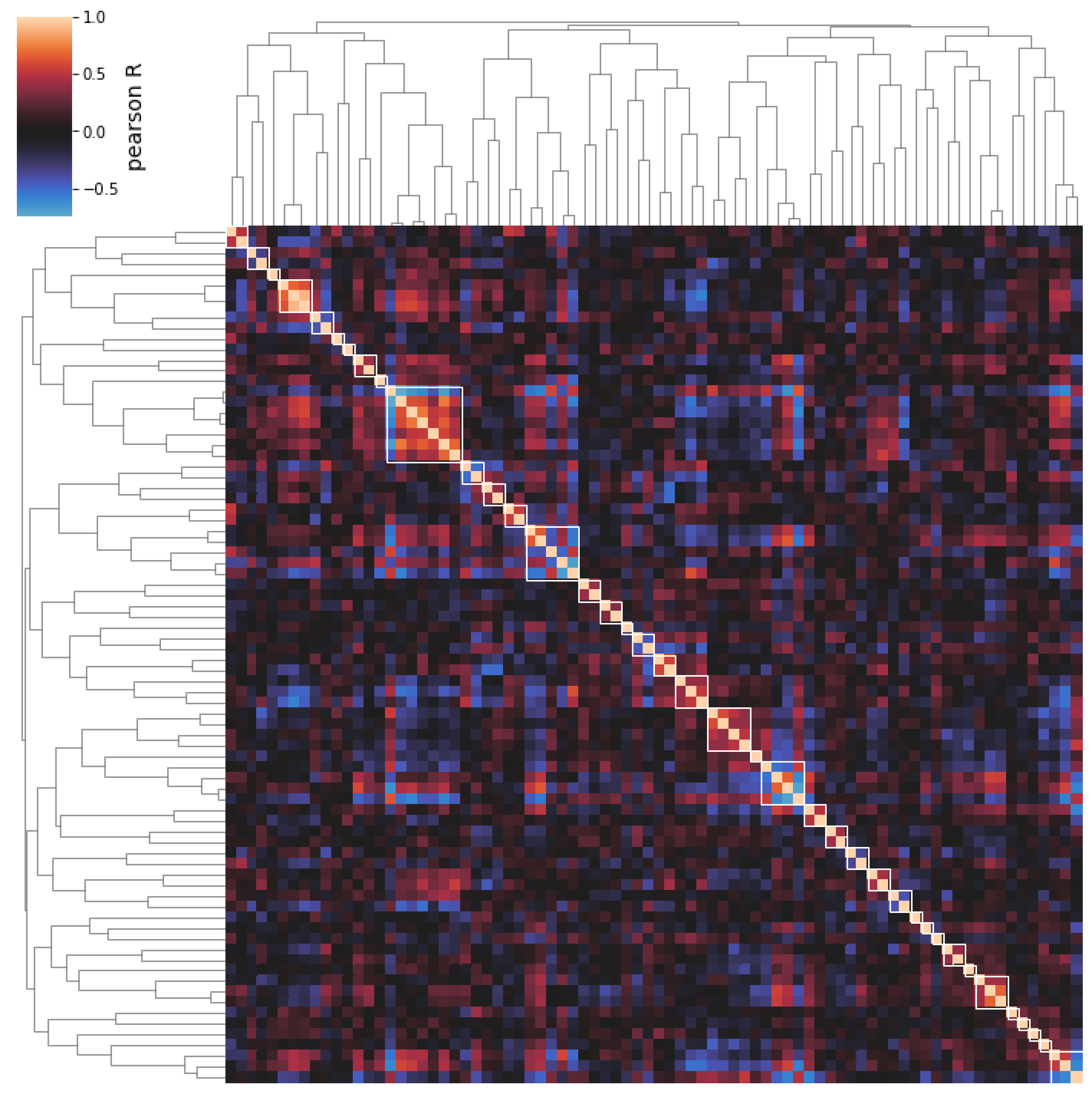

### Supplemental Figure 2 - Core Stress Response Metabolic Map

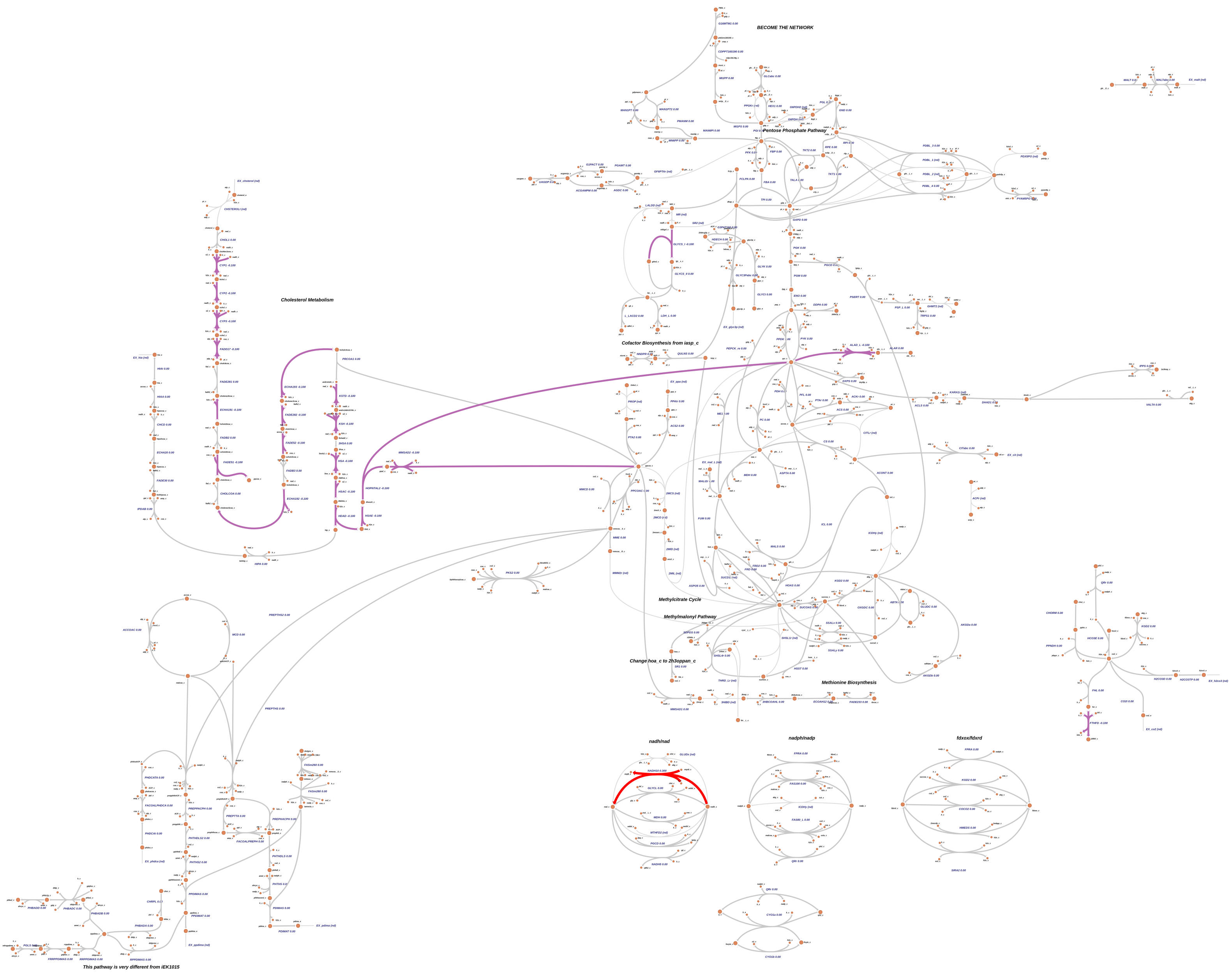
